# Computational Design of Two New 5-HT_1_B Serotonin Receptor Agonists Derived from Naratriptan

**DOI:** 10.64898/2026.05.27.728311

**Authors:** Nicholas L. Martin, Sarah E. Holmes, Justin B. Siegel

## Abstract

Migraine headaches affect over one billion people internationally and can be defined as episodes of acute severe pain wrapping around the head and are normally accompanied by nausea, blurry vision, and sensitivity to light and sound. While triggers that cause migraines may vary among patients, evidence shows they are involved with the trigeminovascular system (a network of blood vessels in the brain in conjunction with the trigeminal nerve). Activation of the trigeminal neurons triggers the release of vasoactive neuropeptides, such as calcitonin gene-related peptide (CGRP), leading to neurogenic inflammation and vasodilation of cranial blood vessels. The development of 5-HT1B serotonin agonist drugs, commonly known as triptans, have been an effective measure of migraine relief. The drugs created in this research were found to have improved docking scores within the 5-HT1B binding site compared to that of naratriptan. The two drugs proposed in this paper would need to undergo further investigation to determine the feasibility of laboratory synthesis and clinical trials.

## INTRODUCTION

Migraines are a common and debilitating neurological disorder characterized by intense headaches often accompanied by nausea, visual disturbances, and increased sensitivity to light and sound.^1^ Affecting over one billion individuals globally, migraines are the third leading cause of disability and can significantly impair daily functioning.^2^ Although precise pathology of migraines is complex and varies from person to person, they are thought to involve trigeminovascular system activation, release of pro-inflammatory neuropeptides, and cause vasoconstriction of cranial vessels.^3^

A primary pharmaceutical approach to treating acute migraine attacks is through the triptan drug class, which is a class of selective (5-HT_1_B/_1_D) receptor agonists.^4^ These drugs act by binding to the selective serotonin receptors along the trigeminal neurons, leading to vasoconstriction of cerebral blood vessels, inhibition of neuropeptide release, and modulation of pain pathways.^5^ Among the several triptans available, avitriptan, eletriptan, naratriptan, and almotriptan were selected to build the library from (Figure 1) due to their favorable pharmacokinetic properties compared to other triptans, however naratriptan was chosen to be the base model of the two candidates based on increased ADMET properties and improved Lipinski’s rules, specifically with the weight of the molecule.^6^

**Figure 1.**
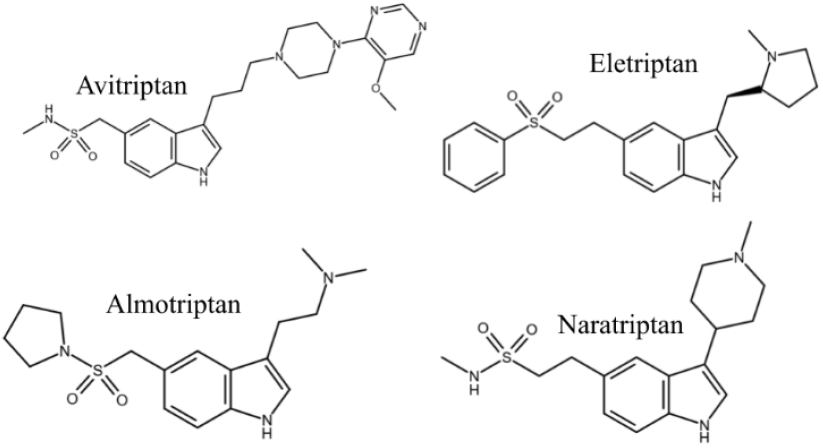
2D Structures of Avitriptan, Eletriptan, Naratriptan, and Almotriptan.

Although the triptan drug class is clinically effective, its activity can lead to off-target vasoconstriction, which may pose risks for certain patient populations, particularly those with hypertension or cardiovascular conditions.^7^ This effect arises from activation of the 5-HT1B receptors on the peripheral vascular system, making it difficult to eliminate alone through selectivity.^8^ The development of new triptan candidates remains important as new structures may enhance central nervous system selectivity. Additionally, enhancing binding affinity may promote more stable ligand receptor interactions, which could contribute to improved pharmacological performance. Overall, structural modification of the triptan scaffold was hypothesized to improve both specificity and binding affinity.

In this paper, two new drug candidates were designed based on naratriptan’s scaffold using a combination of computational methods and chemical intuition. These candidates were evaluated for their predicted binding affinities to the 5-HT_1_B receptor, as well as their pharmacokinetic and toxicological properties, to determine if structural modification could enhance receptor interactions of existing compounds.

## METHODS

The crystal structure of human 5-HT_1_B receptor with bound agonist, donitriptan (PDB ID: 6G79), was obtained from the RCSB Protein Data Bank Website.^9^ All ligand docking simulation is done using OpenEye software.^10^ To design drug candidate 1 based on naratriptan using a bio-isosteric method, the vBrood application^11^ was used to find functional group replacements and sort by ADMET properties.

The naratriptan model was created and optimized in GaussView and Gaussian 09^12^, respectively. Once naratriptan was optimized, Make Receptor was used to define the binding site from the PDB code.^13^ Additionally, VIDA was used to view and analyze drug molecules.^14^ OpenEye OMEGA was used to generate conformer libraries of the drug molecules.^15^ Conformer libraries were docked into the active site using OpenEye FRED, which allows the generation of docking scores.^16^ ADMET properties were scanned using OpenEye Filter^17^, which generates a list of present toxicophores in a spreadsheet. PyMol was used to visualize interactions between ligand and amino acids of the receptors.^18^ BLAST was used to search for homologous proteins;^19^ and the resulting sequences were opened in Jalview^20^ for sequence similarity analyses. Finally, SciFinder^21^ was used to confirm the structures had not already been proposed in literature.^19^

## RESULTS AND DISCUSSION

### Evaluation of Known 5-HT_1_B Agonists

To decide a suitable lead compound for improvement among the four known 5-HT_1_B receptor agonists (avitriptan, eletriptan, naratriptan and almotriptan), each had to be evaluated for their docking scores, ADMET properties, and predicted active site interactions. (Table 1). Among the four, naratriptan had one of the least negative docking scores, indicating the weakest predicted binding affinity to the active site. Despite this, it was chosen for further optimization due to its favorable ADMET properties and modifiable structure.

**Table 1.**
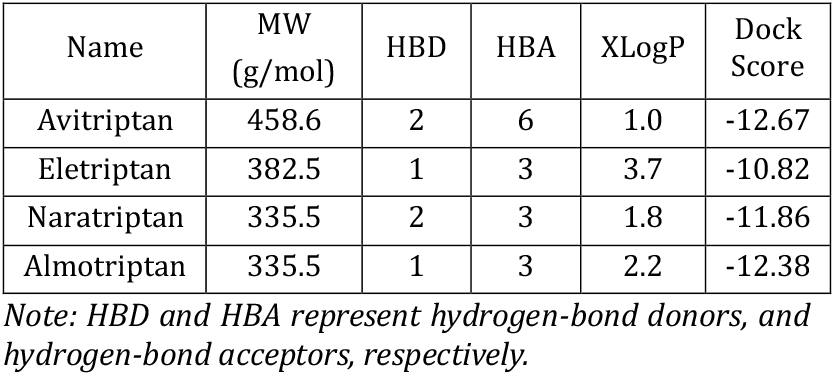
ADMET Properties and Docking Scores of Triptan Library.

All four drugs satisfied Lipinski’s Rule of Five and showed no toxicophores in Filter. Naratriptan tied for lowest molecular weight alongside almotriptan, but naratriptan had a higher donor count and a lower XLogP value, suggesting good solubility and the potential for oral bioavailability. Although its initial binding was weaker, naratriptan’s clean structure and low molecular weight provided larger freedoms to change isosteres, making it a promising candidate for further structure-based designs. Figure 2B,C illustrates the chemical structures of naratriptan, alongside the natural ligand, serotonin, highlighting how this drug mimics serotonin and regulates therapeutic effects. Naratriptan has multiple interactions with amino acids within the active site, including four hydrogen bonding interactions with ASP129 (two 3.1Å, and two 2.4Å bond distances), showing that the interaction is tight.

**Figure 2.**
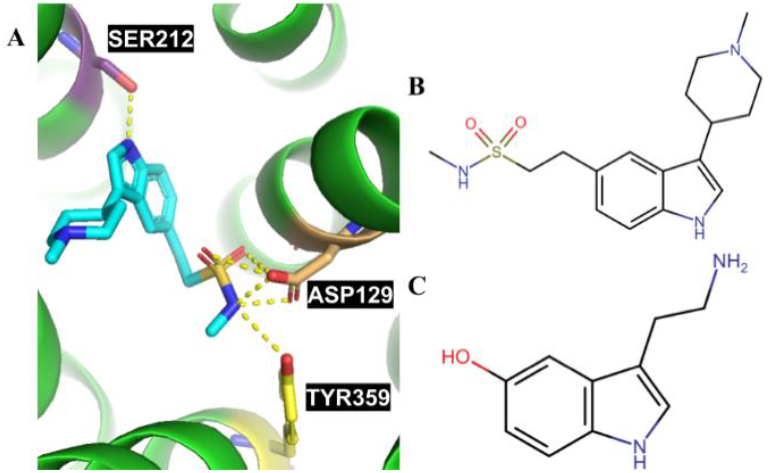
(A) 3D Structure of naratriptan and its interactions with SER212, ASP129, and TYR359. (B) Naratriptan’s 2D structure. (C) The 2D structure for the natural ligand, serotonin (C).

### vBrood Generated 5-HT_1_B Agonist, Candidate 1

Candidate 1 (Figure 3A) was designed computationally by loading naratriptan into vBrood. It was initially run by taking the ring structure on position 3 and replacing it with bioisosteres. This ring was selected based on visualization in Make Receptor, which suggested that the ring’s rigidity may prevent the formation of additional favorable interactions with the active site. After docking the generated compounds, the structure with the highest binding affinity still had the piperidine ring structure that was present in naratriptan, showing that its structure was necessary for an increased binding score. The original structure was then sent through vBrood again to substitute the group at position 5 with various bioisosteres in hopes of achieving a better shape score. The bioisosteres generated were an amide connected to a cyclopropane. This resulted in Generated Candidate 1 as the highest scoring compound (Figure 3), which demonstrated that filling of the pocket was critical for increasing binding.

**Figure 3.**
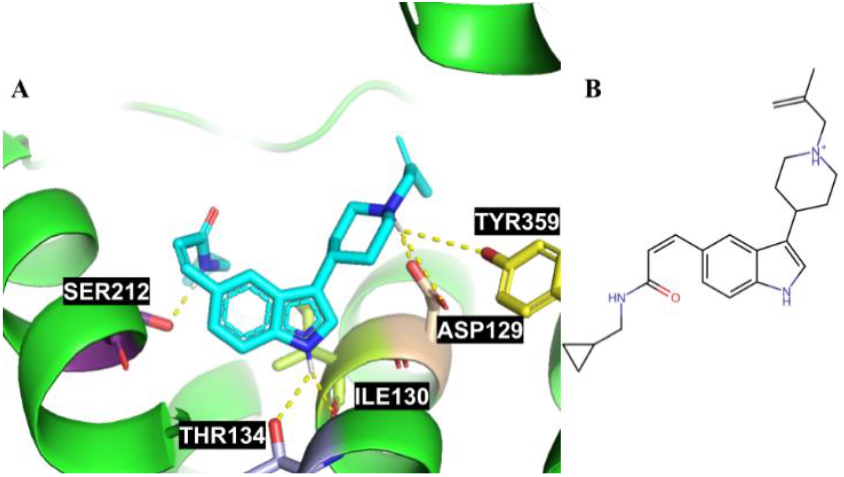
(A) 3D structure of Generated Candidate 1 and its interactions with SER212, THR134, ILE130, ASP129, and TYR359. (B) 2D Structure of Generated Candidate 1.

Candidate 1 had a docking score of -16.11 (Table 2) which is a 35.8% better docking score than naratriptan’s -11.86. It also hydrogen bonds with two additional amino acids, ILE130 (2.2 Å) and THR134 (2.5 Å), resulting in the same amount of hydrogen bonding interactions but spread out over more areas of the active site. These new interactions contributed to the more negative score by -3.64, and the shape filled the pocket making the score more negative by about the same amount. The molecular weight of the drug increased to 378.5 g/mol, hydrogen bond donor count increased by 1, and acceptor count decreased by 2. Its XLogP value did increase significantly to 4.3 but did not violate Lipinski’s Rule of 5. Based on the docking score, molecular interactions, lack of toxicophores, and favorable ADMET properties, Candidate 1 looks to be a promising 5-HT_1_B agonist drug candidate. Candidate 1 is positively charged at physiological pH, which could hinder its ability to cross the blood–brain barrier and limit central nervous system availability. However, established triptan drugs are also charged under physiological conditions and still successfully cross the BBB, likely via an H^+^/organic cation antiporter^22^. Given its structural similarity to these triptans, Candidate 1 may likewise utilize this transport mechanism to achieve BBB permeation and may contribute to central nervous selectivity.

**Table 2.** Naratriptan’s Interactions.

| Residue | Interaction Type | Distance (Å) |
| --- | --- | --- |
| SER212 | Hydrogen Bond | 2.9 |
| ASP129 | Hydrogen Bond | 3.1 |
| ASP129 | Hydrogen Bond | 3.1 |
| ASP129 | Hydrogen Bond | 2.4 |
| ASP129 | Hydrogen Bond | 2.4 |
| TYR359 | Hydrogen Bond | 3.1 |

**Table 3.** Candidate 1’s Interactions.

| Residue | Interaction Type | Distance (Å) |
| --- | --- | --- |
| SER212 | Hydrogen Bond | 1.9 |
| THR134 | Hydrogen Bond | 2.5 |
| ILE130 | Hydrogen Bond | 2.2 |
| ASP129 | Hydrogen Bond | 1.7 |
| ASP129 | Hydrogen Bond | 3.5 |
| TYR359 | Hydrogen Bond | 3.7 |

**Table 4.**
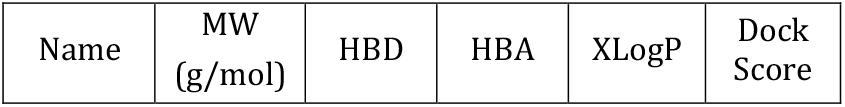
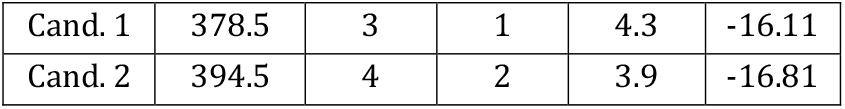
ADMET Properties and Docking Scores of Candidates 1 and 2.

**Table 5.** Candidate 2’s Interactions.

| Residue | Interaction Type | Distance (Å) |
| --- | --- | --- |
| ILE130 | Hydrogen Bond | 2.5 |
| THR134 | Hydrogen Bond | 1.9 |
| ASP129 | Hydrogen Bond | 1.3 |
| ASP129 | Hydrogen Bond | 2.8 |
| CYS133 | Hydrogen Bond | 4.0 |
| TYR359 | Hydrogen Bond | 3.3 |

### Chemical Intuition Driven Design of Candidate 2

Candidate 2 (Figure 4A) was designed based on Candidate 1 in the hopes of fixing some of the problematic ADMET areas. Candidate 2 has a docking score of -16.81 (Table 2) which is a 41.7% increase compared to the original drug, naratriptan.

**Figure 4.**
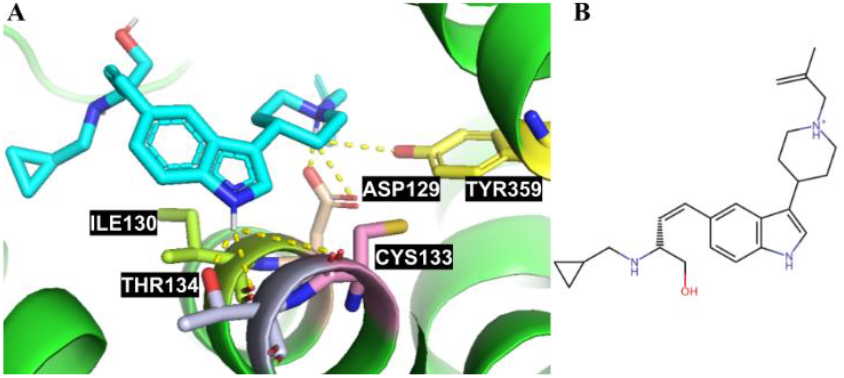
(A) Candidate 2’s Interactions with CYS133, THR134, ILE130, ASP129, and TYR359. (B) 2D Structure of Generated Candidate 2.

Most of the molecular interactions remain the same except that the hydrogen bond with SER212 is no longer present, and a new hydrogen bond with CYS133 is present with a distance of 4.0Å, making it a weaker interaction. THR134 and ASP129 have shorter bond interactions of 1.9Å and 1.3Å, respectively, demonstrating tighter binding than Candidate 1. The molecular weight of Candidate 2 is 394.6 g/mol making it heavier than Candidate 1, but still under the threshold to obey Lipinski’s Rules. Its XLogP value dropped from 4.2 to 3.9 compared to Candidate 1 but is still almost double that of naratriptan. Based on the docking score, molecular interactions, lack of toxicophores, and the other ADMET properties, Candidate 2 is expected to be a better 5-HT_1_B agonist drug than Candidate 1 or naratriptan. Like Candidate 1, it carries a charge, which may limit its ability to cross the blood-brain barrier, but it may also utilize the H^+^/organic cation antiporter like other triptans.

### Off-Target and Preclinical Homology Analysis

To evaluate potential off-target effects of the naratriptan-based drug candidates and their suitability for pre-clinical testing, a protein BLAST and sequence alignment analysis were performed using the 5-HT_1_B receptor FASTA Sequence listed on the RCSB Protein Data Bank. The analysis was carried out across beneficial microbes, pathogenic organisms, and pre-clinical model mammals.

A BLAST^19^ search was conducted against representative beneficial microbes including *Lactobacillus, Bifidobacterium, and Akkermansia muciniphila*. No homologous proteins to the human 5-HT_1_B receptor were identified in any of these species. Since the drug’s target is absent in beneficial microbes, there is no predicted off-target binding, indicating that the two drug candidates are unlikely to disrupt beneficial microbes. This supports the selectivity and safety of the drug candidates in a biological context.

Likewise, no homologous proteins were found in common pathogenic organisms, including *Staphylococcus aureus, Candida albicans, Haemophilus influenzae*, and *Tinea* species. Without a homologous protein in these pathogens, the drug candidates are unlikely to inhibit or interact with their systems involved in infectious diseases. This suggests that the candidates will not contribute to resistance development or unintended microbe targeting.

Highly conserved identity to the human 5-HT_1_B receptor was found in several standard pre-clinical species including *Macaca nemestrina* (pig-tailed macaque), *Canis lupus* (dog), and *Mus musculus* (mouse). A sequence confirmed that the six key drug-binding residues, SER212, ASP129, TYR359, ILE130, THR134, and CYS133, are fully conserved across all standard species. Conservation of the binding suggests that the candidates will likely exhibit similar binding affinities in these models as in humans. These findings support the use of these species in pre-clinical animal testing for evaluating efficacy, safety, and pharmacokinetics of the two candidates.

## CONCLUSION

Migraines affect over one billion people worldwide and remain a major cause of disability, making the development of improved therapies a continuing need. In this study, two novel 5-HT_1_B receptor agonists were designed using naratriptan as the base structure. Candidate 1 showed improved binding affinity and favorable ADMET properties due to added interactions with the active site. Candidate 2 improved upon these results, with the most negative docking score and improved lipophilicity. Both candidates obeyed Lipinski’s Rule of Five, indicating good drug-likeness. Both candidates possess a cationic charge that may limit passive diffusion across the blood–brain barrier but could be moved across by the H^+^/organic cation antiporter, potentially contributing to central nervous system selectivity. Sequence alignment confirmed conservation of the six key binding residues in standard pre-clinical animal models, supporting their suitability for in vivo testing. Further work should focus on synthesis, in vitro testing, and preclinical evaluation to determine their efficacy, safety, and pharmacokinetic behavior.

## AUTHOR INFORMATION

### Present Addresses

† Department of Chemistry, University of California Davis, Davis, California, United States of America, Department of Biochemistry and Molecular Medicine, University of California, Davis, Davis, California, United States of America, Genome Center, University of California Davis, Davis, California, United States of America

### Author Contributions

Research was designed by all authors; all experiments were carried out by N.L.M. The manuscript was written primarily by N.L.M, with contributions from all authors. All authors have given approval to the final version of the manuscript.

## ACKNOWLEDGMENT

Research reported in this publication was supported by UC Davis, the National Science Foundation Award Numbers 1827246, 1805510, 1627539, the National Institute of Environmental Health Sciences of the National Institutes of Health (NIH) under Award Number P42ES004699, UC Davis, NIH Award Number R01 GM 076324-11 and the Rosetta Commons. The content is solely the responsibility of the authors and does not necessarily represent the official views of the National Institutes of Health or National Science Foundation

## REFERENCES

1. Goadsby, P. J.; Holland, P. R.; Martins-Oliveira, M.; Hoffmann, J.; Schankin, C.; Akerman, S. Pathophysiology of Migraine: A Disorder of Sensory Processing. Physiol. Rev. 2017, 97 (2), 553–622.

2. Steiner, T. J.; Stovner, L. J.; Vos, T. GBD 2015: Migraine Is the Third Cause of Disability in under 50s. J. Headache Pain 2016, 17 (1), 104. 10.1186/s10194-016-0699-5

3. Tepper, S. J.; Rapoport, A. M.; Sheftell, F. D. Mechanisms of Action of the 5-HT1B/1D Receptor Agonists. Arch. Neurol. 2002, 59 (7), 1084–1088. 10.1001/archneur.59.7.1084

4. Tfelt-Hansen, P.; De Vries, P.; Saxena, P. R. Triptans in Migraine: A Comparative Review of Pharmacology, Pharmacokinetics and Efficacy. Drugs 2000, 60 (6), 1259–1287. 10.2165/00003495-200060060-00003

5. Goadsby, P. J.; Edvinsson, L. The Trigeminovascular System and Migraine: Studies Characterizing Cerebrovascular and Neuropeptide Changes Seen in Humans and Cats. Ann. Neurol. 1993, 33 (1), 48–56. 10.1002/ana.410330109

6. Havanka, H.; Dahlöf, C.; Pop, P. H.; Diener, H. C.; Winter, P.; Whitehouse, H.; Hassani, H. Efficacy of Naratriptan Tablets in the Acute Treatment of Migraine: A Dose-Ranging Study. Clin. Ther. 2000, 22 (8), 970–980. 10.1016/s0149-2918(00)80068-5

7. Peles, I., Shneyour, R. S., Levanon, E., Steen, Y. M., Abu Salameh, I., Gordon, M., Abuhasira, R., Novack, V., & Ifergane, G. (2025). Cardiovascular risk and triptan usage among patients with migraine. Headache, 65(7), 1095–1106. 10.1111/head.1496

8. Gamoh, S.; Hisa, H.; Yamamoto, R. 5-Hydroxytryptamine Receptors as Targets for Drug Therapies of Vascular-Related Diseases. Biol. Pharm. Bull. 2013, 36 (9), 1410–1415. 10.1248/bpb.b13-00317

9. RCSB Protein Data Bank. PDB ID: 6G79 Coupling specificity of heterotrimeric Go to the serotonin 5-HT1B receptor. https://www.rcsb.org/structure/6G79

10. OEDOCKING, OpenEye Scientific Software: Santa Fe, NM

11. vBROOD, 3.1.6.0; OpenEye Scientific Software: Santa Fe, NM.

12. Gaussian 09; Gaussian, Inc.: Wallingford, CT.

13. Make Receptor, Version 4.1.1.0; OpenEye Scientific Software: Santa Fe, NM.

14. VIDA, Version 5.0.1.0; OpenEye Scientific Software: Santa Fe, NM, 2018.

15. OMEGA 4.1.2.0; OpenEye Scientific Software: Santa Fe, NM.

16. FRED 4.1.1.0; OpenEye Scientific Software: Santa Fe, NM.

17. FILTER; OpenEye Scientific Software: Santa Fe, NM.

18. The PyMOL Molecular Graphics System; Schrödinger, LLC: 2000.

19. BLAST, National Center for Biotechnology Information: Bethesda, MD.

20. Jalview; 2.11.4.1; The Barton Group: University of Dundee, Scotland, UK, 2009.

21. SciFinder 2007.1; Chemical Abstracts Service: Columbus, OH.

22. Svane, N. I.; Pedersen, A. B. V.; Rodenberg, A.; Ozgür, B.; Saaby, L.; Bundgaard, C.; Kristensen, M.; Tfelt-Hansen, P.; Larsen, B. B. The Putative Proton-Coupled Organic Cation Antiporter Is Involved in Uptake of Triptans into Human Brain Capillary Endothelial Cells. Fluids Barriers CNS 2024, 21, 39. https://pmc.ncbi.nlm.nih.gov/articles/PMC11071266/

